# HCMV infection depends on EGLN1-mediated mitochondrial activation to increase dNTP pools for viral DNA replication

**DOI:** 10.1101/2025.10.31.685607

**Authors:** Lucas A Simpson, Diana M Dunn, Wyatt Fales, Zachary J Moore, Jessica H Ciesla, Nicole Waild, Matthew H Raymonda, Isaac S Harris, Joshua Munger

## Abstract

Human cytomegalovirus (HCMV) is a leading cause of congenital infection and morbidity in immunosuppressed populations. Like all viruses, HCMV is an obligate intracellular parasite that extensively remodels host cellular metabolism to support its replication, yet the precise underlying mechanisms and metabolic vulnerabilities remain poorly understood. Using a novel metabolism-focused screening platform, we identified EGLN prolyl hydroxylase activity as critical for HCMV infection. Our studies revealed that HCMV infection depends on EGLN1, which accumulated in mitochondria during infection. Inhibition of EGLN1 expression blocked HCMV-mediated mitochondrial activation, which in turn prevented the production of the dNTP precursors necessary for dNTP pool expansion and viral DNA replication. Further, pharmacological EGLN inhibition attenuated viral infection in a humanized mouse model. Collectively, these data establish EGLN1 as a critical determinant of mitochondrial metabolic remodeling and virally-induced dNTP generation during HCMV infection, highlighting EGLN1 as a promising novel antiviral therapeutic target.

## Introduction

Human cytomegalovirus (HCMV) is a large, double-stranded beta-herpes virus that infects more than 60% of adults worldwide^1^. Following infection, HCMV establishes a latently infected population of cells and persists with the infected host for life. Approximately 8,000 children are born each year with permanent disabilities that arise from congenital CMV infection^2^. Symptoms of congenital HCMV infection can range from mild nonspecific manifestations to severe neurological abnormalities, including microcephaly and death^3^. Additionally, HCMV infection can cause organ transplant rejection, with nearly 40% of infected patients suffering from graft failure^4^. Despite being a major cause of morbidity in immunosuppressed populations and the leading cause of congenital infections in the world, treatment options for HCMV remain limited. Current first-line antivirals suffer from poor oral bioavailability, rapid development of drug resistance, and significant toxicity, including bone marrow suppression, nephrotoxicity, and encephalopathy^5,6^. These limitations underscore the need for additional anti-HCMV therapeutic development.

Viruses rely on cellular metabolic resources to produce viral progeny. HCMV activates numerous metabolic pathways to support infection, including glycolysis^7–9^, glutaminolysis ^10^, and fatty acid biosynthesis^9,11–13^. Additionally, deoxynucleoside triphosphate (dNTP) metabolism has emerged as an important host-pathogen interaction for viruses that have DNA genomes or DNA genome intermediates. For example, cellular SAMHD1 broadly restricts viral infection through its dNTP hydrolase activity, which reduces dNTP pools during inflammatory activation to limit viral DNA replication^14^. To counter this restriction, some viruses target SAMHD1 to maintain elevated dNTP concentrations and support viral DNA synthesis^15,16^. Virally-associated nucleotide metabolism is also frequently targeted therapeutically for treating hepatitis B, HIV, cytomegalovirus, and herpes simplex virus infections^17–20^. However, despite their importance for infection and therapeutic relevance, most virally-associated metabolic dependencies remain unknown, along with the specific mechanisms through which viruses modulate cellular metabolic controls.

In this study, we employed a metabolism-focused screening platform to identify HCMV-associated metabolic vulnerabilities. Several inhibitors of amino acid, nucleotide, and lipid metabolism attenuated HCMV infection. Furthermore, pharmacological inhibition of the EGLN family of prolyl hydroxylases substantially reduced viral replication. Among the three EGLN family members, EGLN1 was the dominant enzyme contributing to successful infection, proving essential for HCMV activation of TCA cycle anaplerosis and stimulation of mitochondrial metabolism and biogenesis. Additionally, EGLN1-mediated increases in mitochondrial activity are essential for HCMV-induced deoxynucleotide production, which is critical for viral DNA synthesis and infectious progeny production. Our findings identify EGLN1 as a major regulator of mitochondrial function during viral infection.

## Results

### Metabolism focused pharmacological screening identifies inhibitors of HCMV infection

To uncover metabolic vulnerabilities associated with HCMV infection, we designed a screening platform utilizing a compound library targeting metabolism-associated enzymes. Each compound was arrayed over a 10-point dose curve ranging from 20 μM to 1 nM^21,22^. Following multiple rounds of replication, the impact of these compounds was assessed by quantifying HCMV-encoded GFP expression using an imaging-based readout (Figure 1A). Optimization of screening conditions yielded a Z-factor of 0.95 (Supplemental Figure 1), indicating a robust screening assay²³. In addition to monitoring GFP infected cells, a parallel screen monitored the impact of compound treatment on mock-infected cells, enabling comparison of compound doses that inhibit viral infection versus those that are cytotoxic to uninfected cells.

**Figure 1.**
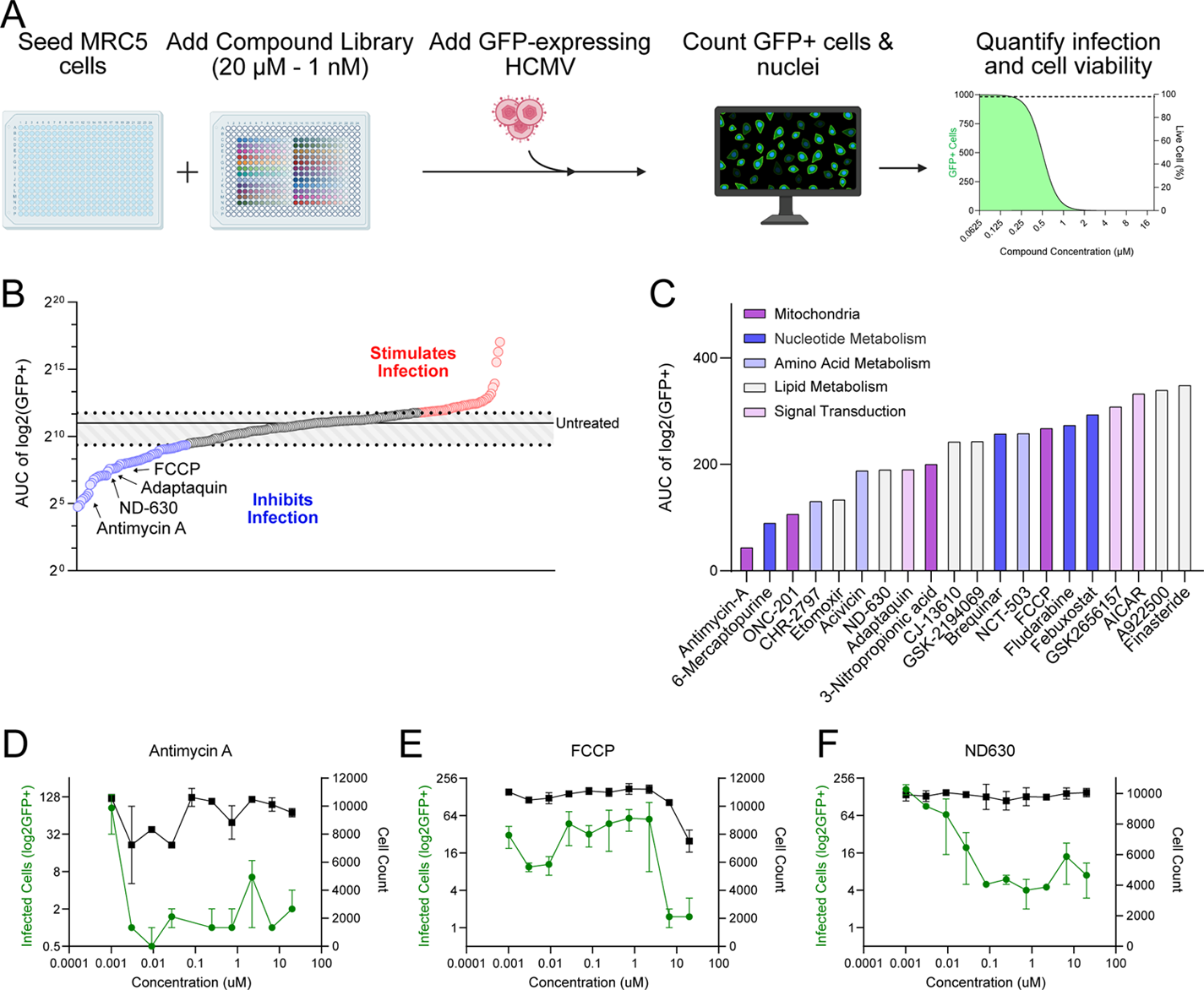
A high-throughput metabolic inhibitor screen identifies multiple inhibitors of HCMV infection. (A) Schematic representing high-throughput screening method to identify metabolic vulnerabilities of HCMV. MRC5-hT fibroblasts were treated with our compound library and then infected with HCMV-GFP (AD169, MOI = 0.01). GFP-expressing cells and nuclei were quantified 9 days post infection. Hit compounds were determined by comparing the AUC of the GFP-expressing cell counts relative to untreated infected cells. (B) Area under the curve (AUC) of infected GFP-expressing cell counts were quantified (N=2) (C) Pathways associated with the top 20 compounds, excluding compounds with cytotoxic effects. (D-F) Screening results for select compounds. GFP-positive infected cells (green) and mock infected cell number (black) are represented (Mean ± SEM, N=2).

Several compounds inhibited infection, while others appeared to stimulate infection, as determined based on the areas under the infected cell dose response curves compared to untreated infections (Figure 1B, Supplemental File 1). Multiple inhibitors targeted metabolic pathways previously reported to be important for infection, including lipid^9,11^, nucleotide^23^, and amino acid metabolism^10^, as well as mitochondrial electron transport^24,25^. Among the top 20 compound hits, six targeted lipid metabolism, four targeted nucleotide metabolism, four targeted mitochondrial processes, and three targeted amino acid metabolism (Figure 1C, Supplemental File 1).

Antimycin-A and FCCP, which target mitochondrial electron transport, both limited viral infection, although Antimycin-A did so at lower concentrations (Figure 1D &1E). ND630, which inhibits lipid synthesis through inhibition of acetyl-CoA carboxylase (ACC), attenuated infection with negligible impact on uninfected cells (Figure 1F). These findings are consistent with previous reports analyzing the impact of alternative ACC inhibitors on HCMV infection^9,11^. Collectively, these results reinforce the importance of these metabolic pathways for viral infection, while identifying potential new compounds with antiviral activity.

### Adaptaquin treatment attenuates the production of HCMV progeny

Adaptaquin was one of the top inhibitors identified in the screen (Figure 1C). Adaptaquin inhibits the EGLN family of prolyl hydroxylases^26,27^, enzymes well known for their regulation of HIF1α levels through oxygen-dependent hydroxylation of specific proline residues in HIF1α, leading to its degradation^28–30^. Subsequent validation experiments confirmed that adapataquin inhibited HCMV viral spread (Figure 2A) at concentrations that did not impact cellular viability (Figure 2B&C). Adaptaquin treatment greatly reduced the production of viral progeny to the limit of detection at either low or high MOI infection (Figure 2D&E) and blocked the viral spread of a limited passage, more clinically relevant strain of HCMV (TB40/E) (Figure 2F&G).

**Figure 2.**
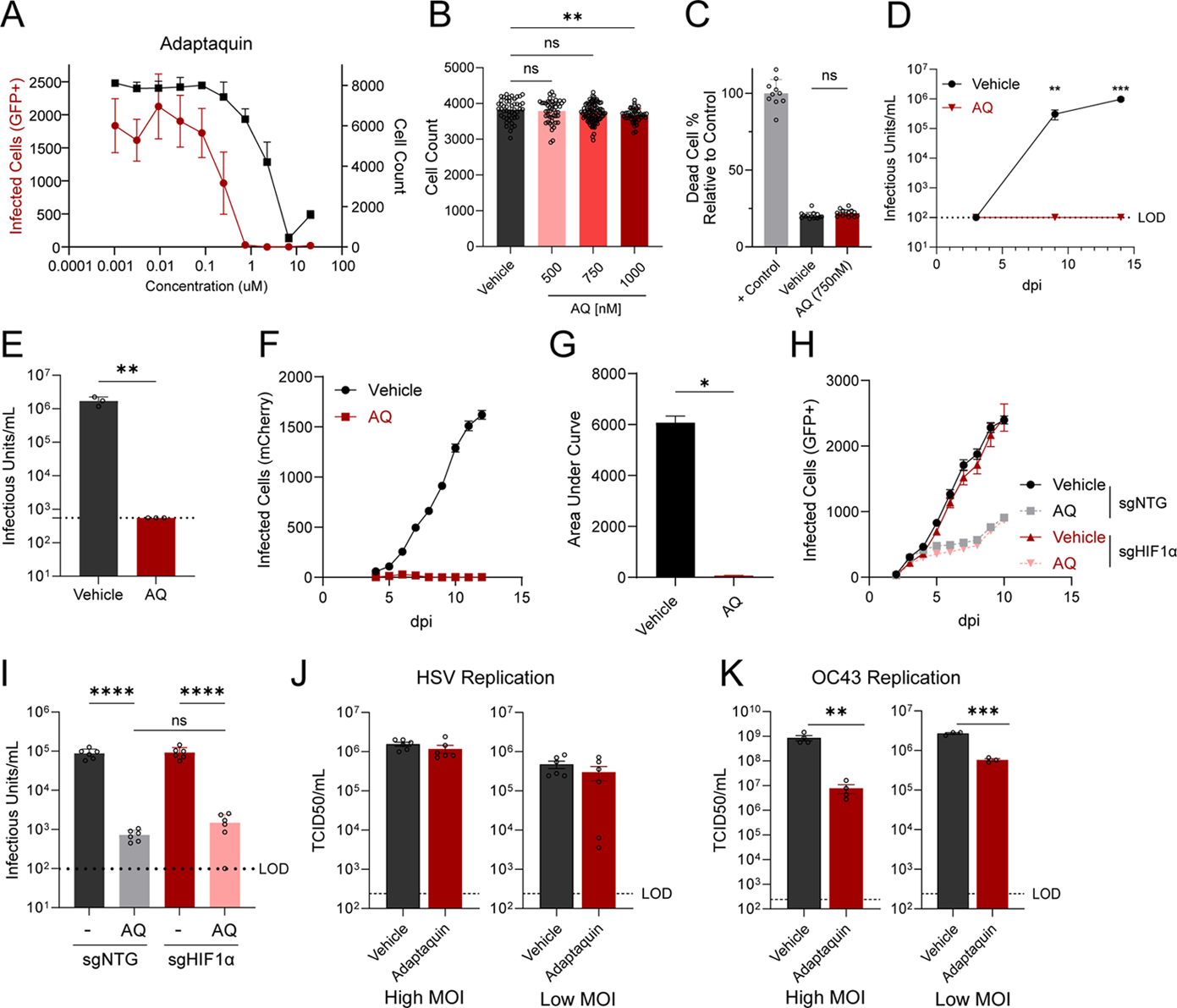
Adaptaquin inhibits HCMV infection. (A) Dose response of cells were treated with adaptaquin (AQ) and infected with HCMV-GFP (AD169, MOI = 0.01). GFP-positive infected cells (red) and mock infected cell number (black) are represented (Mean, N=4, +/- SE). (B) Cell counts of uninfected fibroblasts treated with AQ for 5 d that were fixed and stained with Hoescht (Mean ± SEM, N=48). (C) Dead cell percentages of uninfected fibroblasts treated with AQ (750 nM) for 5 d that were fixed and stained with propidium iodine, normalized to positive control methanol fixed cells (Mean ± SEM, N=24). (D) Viral titer of fibroblasts infected with HCMV-GFP (AD169, MOI = 0.01) and treated with AQ (750 nM). Progeny production was assessed at 3, 9, and 14 d post infection (Mean ± SEM, N=3). (E) Viral titer of fibroblasts infected with HCMV-GFP (AD169, MOI = 3) and treated with AQ (750 nM). Progeny production was assessed at 5 d post infection (Mean ± SEM, N=3). (F-G) Viral spread and area under the curve of fibroblasts infected with HCMV-mCherry (TB40/E, MOI = 0.01) and treated with AQ (750 nM) (Mean ± SEM, N=5). (H) Viral spread of fibroblasts lacking HIF1α expression and non-targeting (NTG) control cells infected with HCMV-GFP (AD169, MOI = 0.01) and treated with AQ (750 nM) (Mean ± SEM, N=10). (I) Viral titer of fibroblasts lacking HIF1α expression and NTG control cells infected with HCMV-GFP (AD169, MOI = 3) and treated with AQ (750 nM). Progeny production was assessed at 5 d post infection (Mean ± SEM, N=6). (J) Viral titer of fibroblasts infected with HSV (MOI = 5, MOI = 0.05, respectively) and treated with AQ (750 nM). Progeny production was assessed 24 hours post infection (hpi) (Mean ± SEM, N=6). (K) Viral titer of fibroblasts infected with OC43 (MOI = 1, MOI = 0.05, respectively) and treated with AQ (750 nM). Progeny production was assessed 24 hpi (Mean ± SEM, N=4, 3, respectively). The statistical significance is displayed as follows: *p < 0.05; **p < 0.01; ***p < 0.001; ****p < 0.0001; ns, not significant.

Given that EGLN inhibition would be predicted to increase HIF1α levels^28–30^, we investigated whether adaptaquin-mediated inhibition of infection depends on HIF1α. We utilized fibroblasts in which HIF1α had been knocked out using CRISPR-Cas9 (Supplemental Figure 2)^31^. Adaptaquin treatment caused a similar reduction in viral spread in HIF1α knockout versus control cells (Figure 2H). Viral titers produced in the presence of adaptaquin were also similar between control and the HIF1α knockout cells (Figure 2I). These data indicate that adaptaquin-mediated inhibition of HCMV replication is independent of HIF1α.

To determine whether adaptaquin’s inhibition of viral replication is specific to HCMV, we assessed the impact of adaptaquin treatment on replication of herpes simplex virus (HSV) and the OC43 coronavirus. Adaptaquin treatment did not impact HSV replication under high or low MOI conditions (Figure 2J), while OC43 replication was significantly reduced (Figure 2K).

These data suggest that adaptaquin’s inhibition of viral replication is not limited to HCMV, but that adaptaquin does not inhibit all viruses non-specifically, which might be expected if cytotoxicity were a contributing factor.

### Adaptaquin attenuates HCMV infection at a late stage, blocking viral DNA replication

HCMV gene expression occurs in tightly regulated sequential steps during infection, with viral genes classified as immediate-early (IE), early (E), and late (L)^32^. IE gene expression does not require de novo viral gene expression upon infection, whereas E gene expression requires immediate-early gene expression (Figure 3A). Late gene expression depends on viral DNA replication (Figure 3A). To examine how adaptaquin treatment impacts the HCMV life cycle, we analyzed viral gene expression via proteomics at 24 and 72 hours post-infection (hpi) (Supplemental File 2).

**Figure 3.**
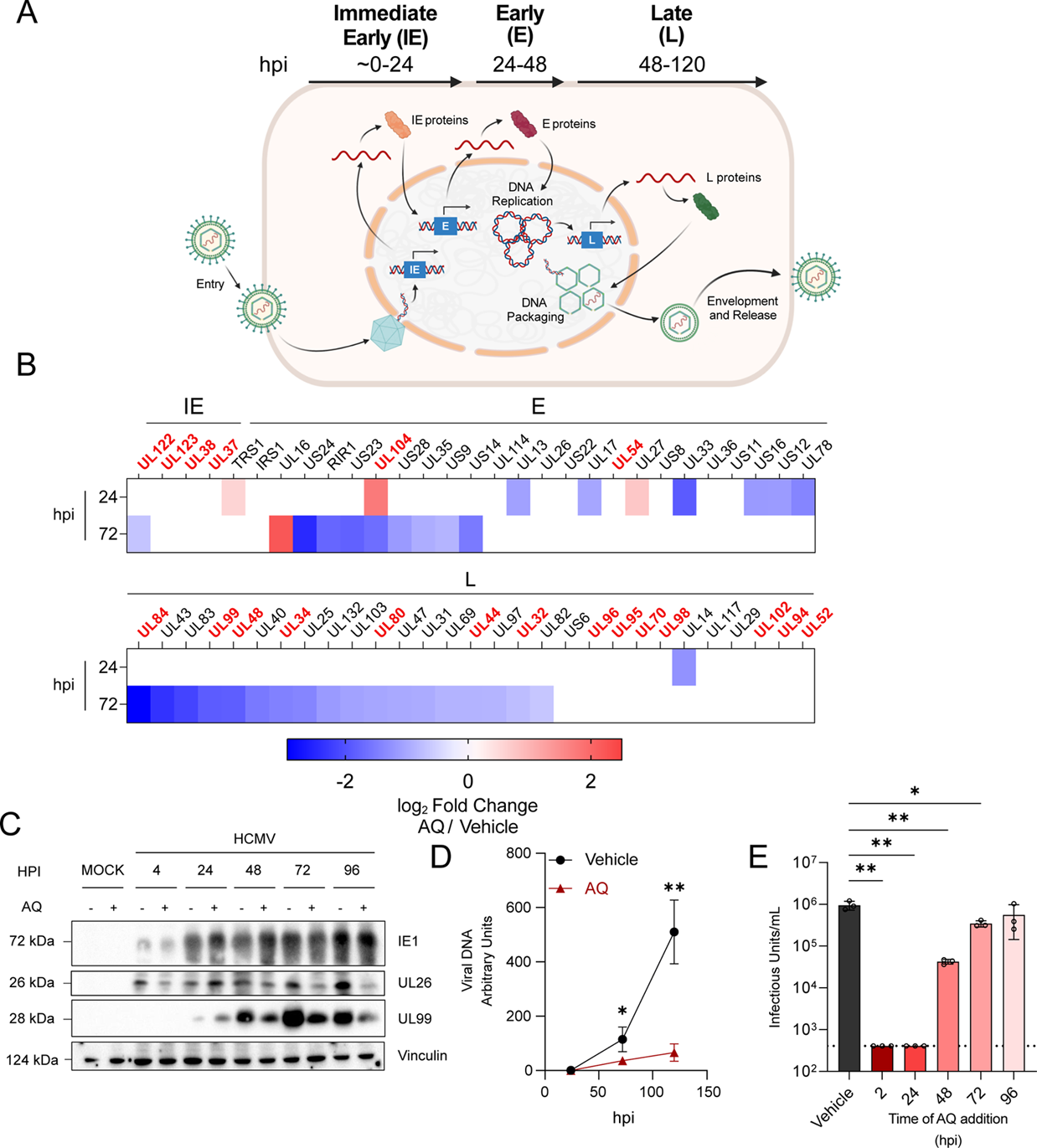
Adaptaquin treatment inhibits viral DNA synthesis and late protein production. (A) Schematic representing the viral life cycle of HCMV. (B) Viral protein abundances determined by LC-MS/MS from infected fibroblasts (MOI = 1.5) and treated with adaptaquin (AQ, 750 nM). Protein samples were obtained 24 and 72 hpi, respectively. Fold change was calculated comparing AQ vs Vehicle treated samples. Proteins labeled in red are essential for HCMV replication (Mean, N=3). (C) Immunoblot of infected fibroblast cell lysates (MOI = 3) and treated with AQ (750 nM), harvested at designated time points. (D) Viral DNA quantification of infected fibroblasts (MOI = 1.5) treated with AQ (750 nM) using genomic DNA harvested at 24, 72, and 120 hpi and quantitative PCR using primers for IE1 (Mean ± SEM, N=3). (E) Viral titer of infected fibroblasts (MOI = 3) and treated with AQ (750 nM) at designated time points. Progeny production was assessed 5 dpi (Mean ± SEM, N=3). The statistical significance is displayed as follows: *p < 0.05; **p < 0.01; ***p < 0.001; ****p < 0.0001; ns, not significant.

At 24 hpi, adaptaquin treatment reduced the accumulation of select non-essential early proteins, but did not significantly impact the accumulation of immediate early proteins (Figure 3B). The largest adaptaquin-induced reductions occurred in L proteins (Figure 3B), many of which are essential for viral infection (highlighted in red)^33,34^. Western blot analysis of individual representatives of these kinetic classes yielded similar results. At earlier infection timepoints, accumulation of immediate-early 1 protein (IE1) and the early UL26 protein were not substantially affected at 24 hpi (Figure 3C), whereas protein levels of the late gene UL99 (pp28) were substantially reduced at 48 hpi and later timepoints (Figure 3C). Adaptaquin’s inhibition of late protein accumulation suggested it might inhibit viral DNA synthesis, which is required for late gene expression. Consistent with this hypothesis, adaptaquin treatment substantially reduced viral DNA accumulation relative to controls (Figure 3D).

Adaptaquin’s inhibition of viral DNA synthesis suggested that it might be capable of inhibiting viral infection at the later stages of the viral life cycle. We therefore tested adaptaquin’s ability to inhibit productive infection when added at different time points. Upon addition at 24 hpi, a time when viral DNA replication is initiating, adaptaquin reduced the production of viral progeny to the limit of detection (Figure 3E). When added at 48 hpi, adaptaquin treatment reduced viral replication by >10-fold and over 2-fold when added at 72 hpi (Figure 3E). These data suggest that adaptaquin is capable of inhibiting viral replication after infection is established and indicate that adaptaquin inhibits a late stage of infection, substantially attenuating viral DNA replication and the subsequent expression of essential late viral proteins.

### HCMV viral DNA synthesis depends on EGLN1 and EGLN2

Adaptaquin inhibits the EGLN family of prolyl hydroxylases ^26,27^. HCMV infection moderately induced EGLN1 and EGLN2 RNA accumulation at specific time points (Figure 4A&B). EGLN3 RNA expression was much more dramatically induced, with a 10-fold or greater induction throughout infection (Figure 4C), consistent with reports that EGLN3 expression can be induced by inflammatory signaling ^35,36^, which is activated during HCMV infection^37^. Given adaptaquin’s anti-HCMV activity, we tested whether inhibition of EGLN gene expression would attenuate HCMV replication. CRISPRi knockdown of EGLN1 substantially reduced its expression (Figure 4D) and strongly attenuated HCMV viral spread (Figure 4 E&F). Similar knockdown of EGLN2 reduced its expression (Figure 4G), and reduced viral spread, albeit to a lesser extent than EGLN1 (Figure 4H&I).

**Figure 4.**
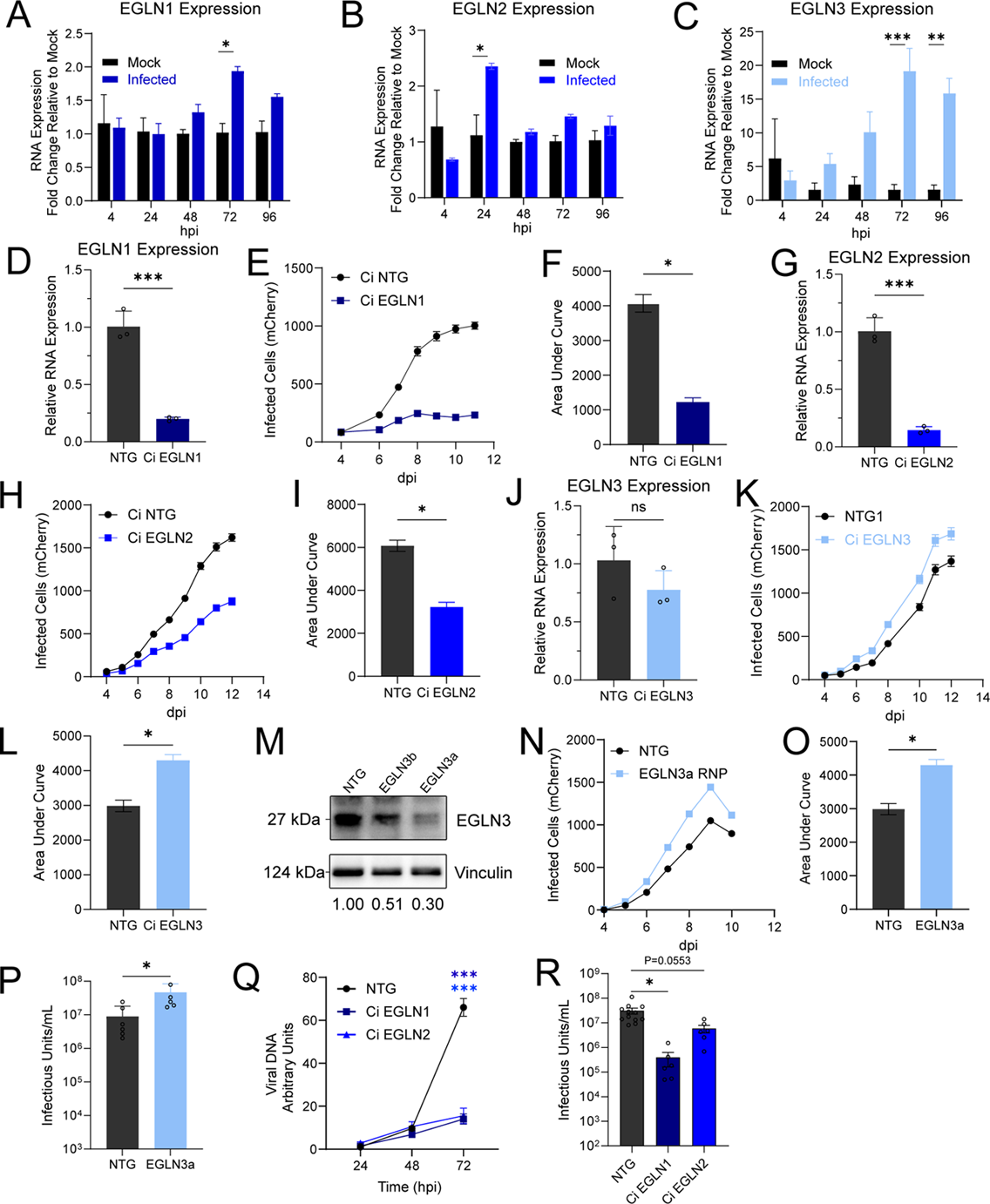
CRISPR-mediated inhibition of EGLN1 and EGLN2 expression inhibits HCMV replication. (A-C) EGLN expression in infected fibroblasts as determined by qPCR at designated time points (MOI=1.5, Mean ± SEM, N=3). (D) EGLN1 expression in fibroblasts with CRISPRi-targeted EGLN1 as determined by qPCR (Mean ± SEM, N=3). (E-F) Viral spread and area under the curve of CRISPRi EGLN1 knockdown cells infected with HCMV-mCherry (TB40/E, MOI = 0.01, Mean ± SEM, N=20). (G) EGLN2 expression in fibroblasts with CRISPRi-targeted EGLN2 as determined by qPCR (Mean ± SEM, N=3). (H-I) Viral spread and area under the curve of CRISPRi EGLN2 knockdown cells infected with HCMV-mCherry (TB40/E, MOI = 0.01, Mean ± SEM, N=20). (J) EGLN3 expression in fibroblasts with CRISPRi-targeted EGLN3 as determined by qPCR (Mean ± SEM, N=3). (K-L) Viral spread and area under the curve of EGLN3 knockdown cells infected with HCMV-mCherry (TB40/E, MOI=0.01, Mean ± SEM, N=20). (M) Immunoblot of cell lysates from fibroblasts transfected with an RNP targeting EGLN3 and a NTG. Protein was harvested following cell recovery and assessed for EGLN3 expression via Western blotting. Normalized EGLN1 abundances represented below the blot. (N-O) Viral spread and area under the curve of EGLN3 RNP cells infected with HCMV-mCherry (TB40/E, MOI = 0.01, Mean ± SEM, N=20). (P) Viral titer of EGLN3a RNP cells infected with HCMV-GFP (AD169, MOI = 3, 5 dpi, Mean ± SEM, N=3). (Q) Viral DNA quantification of EGLN1 and EGLN2 knockdown cells infected with HCMV-GFP from genomic DNA harvested at designated time points by quantitative PCR using primers for IE1 (AD169, Mean ± SEM, N=3). (R) Viral titer of CRISPRi EGLN1 and EGLN2 cells infected with HCMV-GFP (AD169, MOI = 3). Progeny production was assessed 5 dpi (Mean ± SEM, N=3). The statistical significance is displayed as follows: *p < 0.05; **p < 0.01; ***p < 0.001; ****p < 0.0001; ns, not significant.

CRISPRi knockdown EGLN3 was less effective at reducing EGLN3 RNA levels (Figure 4J), although it modestly increased HCMV replication (Figure 4K&L). To increase the EGLN3 knockdown efficiency, we utilized CRISPRn ribonucleoproteins (RNP). CRISPRn RNP knockout reduced EGLN3 protein levels by ∼70% (Figure 4M) and increased both HCMV viral spread and viral titers (Figure 4N-P), similar to what was observed with CRISPRi knockdown of EGLN3. These results suggest that EGLN3 expression modestly restricts HCMV infection.

To further expand on the attenuation of HCMV replication following knockdown of EGLN1 and EGLN2, we next assessed viral DNA accumulation and the production of infectious virions. Similar to adaptaquin treatment, EGLN1 or EGLN2 knockdown inhibited the accumulation of HCMV viral DNA (Figure 4Q). EGLN1 knockdown reduced the production of infectious virions by >50-fold, whereas EGLN2 knockdown reduced viral production by ∼5-fold (Figure 4R). Collectively, these results demonstrate that EGLN1 and EGLN2 are important for HCMV viral infection.

Notably, expression of the various EGLN genes was responsive to targeting other family members. For example, targeting EGLN1 or EGLN3 induced EGLN2, whereas targeting EGLN2 induced EGLN1 (Supplemental Figure 3). HIF1α regulates the transcription of EGLN family members^38^ and given that HIF1α is in turn regulated by EGLN activity, it could be expected that reduced expression of a specific EGLN would impact the expression of other EGLN family members. This intertwined regulation of EGLN expression complicates straightforward interpretations with respect to their contributions to infection.

### EGLN1 is necessary for HCMV-induced mitochondrial activation

In breast cancer cells, EGLN1 reportedly translocates to the mitochondria independently of HIF1α^39,40^. We therefore examined EGLN contributions to mitochondrial localization and function during HCMV infection. Consistent with previous findings^41,42^, HCMV infection increased basal and maximal respiration (Figure 5A, Supplemental Figure 4). However, this activation was blocked by adaptaquin treatment or EGLN1 knockdown (Figure 5A-B). In uninfected cells, EGLN1 inhibition modestly inhibited uncoupled and basal respiration (Figure 5A-B). In contrast to EGLN1, EGLN2 knockdown induced a slight increase in the maximal respiration of infected cells (Figure 5C), suggesting EGLN1 is an important determinant for HCMV’s induction of mitochondrial respiration. Consistent with EGLN1 being important for HCMV-induced mitochondrial activation, infection induced the accumulation of EGLN1 in mitochondrial fractions (Figure 5D).

**Figure 5.**
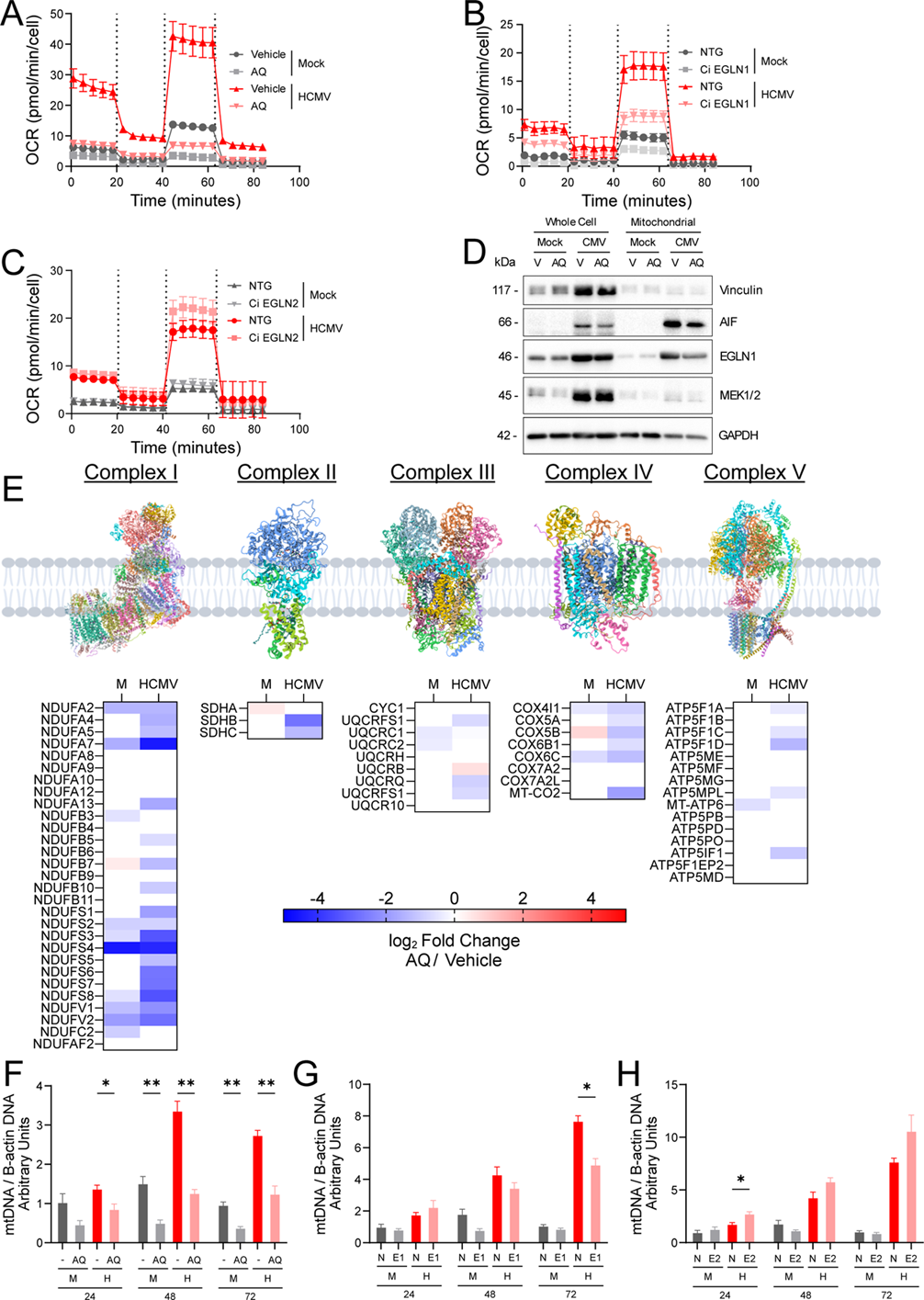
EGLN1 is necessary for HCMV-mediated activation of respiration and mitochondrial biogenesis. (A-C) Oxygen consumption for adaptaquin (AQ, 750 nM) treated (A), EGLN1 knockdown (B), and EGLN2 knockdown cells (C). Oxygen consumption normalized to cell count (MOI=1.5, Mean ± SEM, N=20). (D) Immunoblot of cell lysates from infected fibroblasts (MOI = 1.5) treated with AQ (750 nM). 2% cell equivalents of whole cell or mitochondrial fractions harvested 48 hpi were loaded for subsequent western blot analysis (N=3, independent experiments). (E) A schematic of the electron transport chain proteins and their quantified subunits that were determined as described in Figure 3B (72 hpi, MOI=1.5). Fold change was calculated comparing AQ vs Vehicle treated samples. (F-H) mtDNA quantity from AQ treated (A), EGLN1 knockdown (B), and EGLN2 knockdown cells (C). Genomic DNA was harvested from infected cells (MOI = 1.5) at designated time points, using primers for mitochondrial ND1 and B-actin (Mean ± SEM, N=3). The statistical significance is displayed as follows: *p < 0.05; **p < 0.01; ***p < 0.001; ****p < 0.0001; ns, not significant.

Proteomic analysis indicated that adaptaquin treatment reduced key proteins involved in mitochondrial metabolism and physiology, including the TCA cycle and Complex I biogenesis (Supplemental File 3). Adaptaquin treatment significantly decreased HCMV-induced accumulation of various respiratory chain subunits (Figure 5E). Given that HCMV is known to induce mitochondrial biogenesis in infected cells^41–43^, we examined HCMV-induced activation of mitochondrial DNA replication. As previously observed, HCMV infection induced mtDNA accumulation, but this induction was attenuated by adaptaquin treatment or CRISPRi-mediated targeting of EGLN1, but not EGLN2 (Figure 5F-H). Collectively, these results indicate that EGLN1 expression and activity are important for HCMV-mediated activation of respiration and mitochondrial biogenesis.

### EGLN activity is necessary for HCMV-mediated induction of TCA cycle-related metabolites

To more globally examine how EGLN activity contributes to HCMV-mediated metabolic remodeling, we utilized LC-MS/MS to assess the impact of EGLN inhibition on metabolite levels (Supplemental File 4-6). Principal component analysis (PCA) of the resulting data indicated that adaptaquin treatment substantially impacted virally-induced metabolic remodeling (Figure 6A). Infection induced the largest shifts in the data, regardless of treatment, as shown by the shifts in principal component 1 (PC1) (Figure 6A). Adaptaquin-treated HCMV-infected cells were largely separated from infected control cells along PC2 (Figure 6A), suggesting that adaptaquin substantially impacts HCMV-mediated modulation of cellular metabolism. In contrast, adaptaquin-treated mock-infected cells exhibited significant overlap with control-treated mock-infected cells (Figure 6A), suggesting that adaptaquin’s metabolic effects are largely infection-specific.

**Figure 6.**
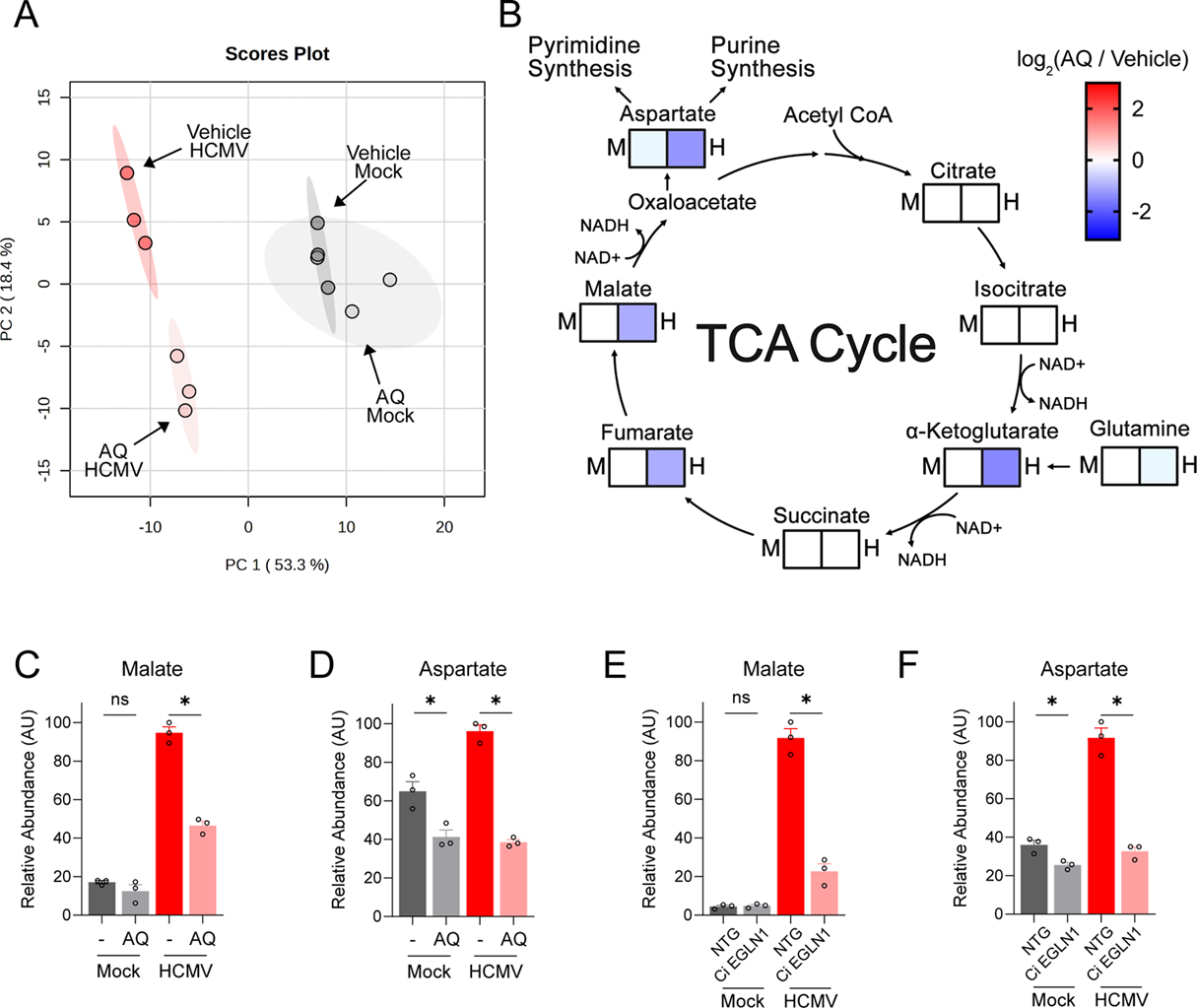
EGLN1 is necessary for HCMV-mediated activation of TCA cycle anaplerosis and the induction of aspartate concentrations. (A) Principal component analysis of quantified metabolites from metabolomics performed in infected fibroblasts (MOI = 1.5) treated with adaptaquin (AQ, 750 nM) and harvested 48 hpi. Abundances were quantified via LC-MS/MS (Mean ± SEM, N=3). (B) A schematic representing the significantly impacted metabolites involved in the TCA cycle for mock (M) and infected (H) cells. Metabolite fold changes were determined comparing AQ treatment vs vehicle treatment. Non-significant changes are displayed in white. (C-D) Relative abundance of malate and aspartate for AQ treatment (750 nM), respectively. (Mean ± SEM, N=3) (E-F) Relative abundance of malate and aspartate in control and CRISPRi EGLN1 knockdown cells. (Mean ± SEM, N=3). The statistical significance is displayed as follows: *p < 0.05 by one-way ANOVA analysis with FDR correction.

The metabolites most impacted by adaptaquin treatment during infection included various TCA cycle and TCA-adjacent compounds (Figure 6B-D, Supplemental File 4). These included glutamine, α-ketoglutarate, fumarate, malate, and aspartate, all of which were significantly decreased by adaptaquin treatment during infection, but were not substantially impacted in mock-infected cells (Figure 6C&D). Similar results were observed with CRISPRi knockdown EGLN1. Malate and Aspartate were substantially reduced in HCMV-infected cells relative to non-targeting controls, and this effect was largely specific for viral infection (Figure 6E&F). These data indicate that, in addition to attenuating HCMV-activated mitochondrial respiration, EGLN1 is important for HCMV-mediated induction of TCA cycle metabolite pools.

### EGLN1-dependent aspartate generation is critical for HCMV-induced viral DNA replication

TCA cycle-linked aspartate production is critical for both pyrimidine and purine biosynthesis, as aspartate provides nitrogen for purine ring biosynthesis, and carbon and nitrogen for the production of pyrimidines (Figure 7A)^44^. The observed reductions in aspartate levels upon EGLN inhibition (Figure 6D&F) could therefore result in reduced nucleotide biosynthesis and the attenuation of HCMV DNA replication. Consistent with this hypothesis, adaptaquin treatment or CRISPRi-mediated knockdown of EGLN1 substantially attenuated dNTP accumulation during HCMV infection (Figure 7B&C).

**Figure 7.**
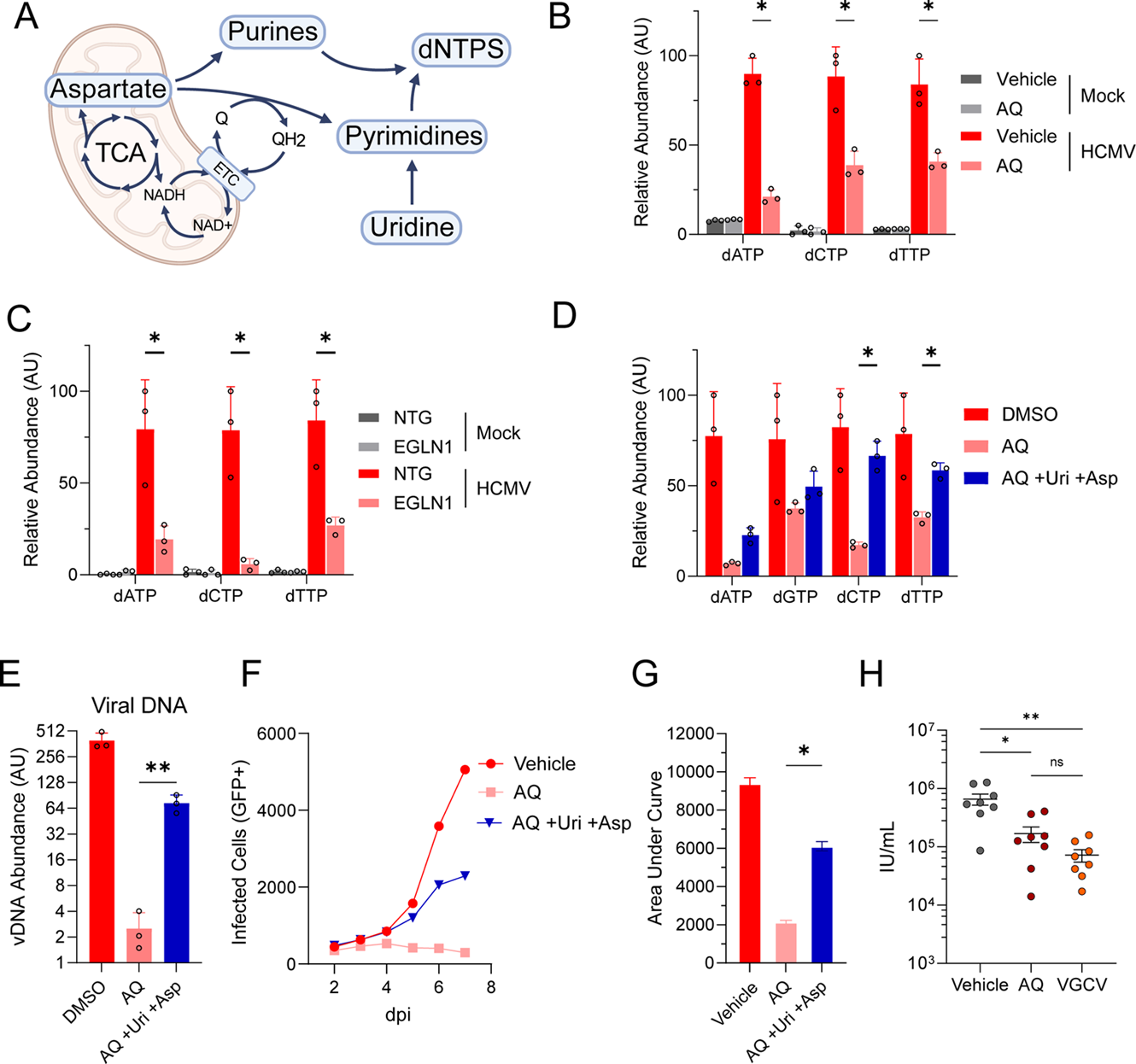
EGLN1-mediated mitochondrial activation provides the nucleotide precursors necessary to support dNTP pool expansion and drive viral DNA replication. (A) Schematic representing the links between the electron transport chain (ETC), NAD+, aspartate, uridine, and nucleotide synthesis. (B-D) LC-MS/MS based quantification of dNTPs in control and adaptaquin (AQ, 750 nM) treated (B), EGLN1 knockdown (C) and AQ treated fibroblasts supplemented with 20 mM aspartate and 0.41 mM uridine (D). Metabolite abundances were quantified as described in Figure 5A (48 hpi, MOI=1.5, Mean ± SEM, N=3). (E) Viral DNA quantification of infected fibroblasts (MOI = 1.5) using viral genomic DNA harvested at 72 hpi (Mean ± SEM, N=3). (F-G) Viral spread and area under the curve of infected fibroblasts (MOI = 0.01) treated with AQ (750 nM) and supplemented with aspartate and uridine (MOI=0.01, Mean ± SEM, N=12). (H) Viral titers of matrigel plugs following antiviral treatment *in* vivo. MRC5-hT cells were infected with HCMV-mCherry (TB40/E, MOI = 0.02) and incubated for 24 hours, prior to seeding (1.6 x 10^6^ cells per injection) into matrigel suspension. Mice were treated with adaptaquin (50 mg/kg, i.p., daily) for 9 days post implantation, prior to plug removal and viral quantification. Titer values are normalized by gram of harvested matrigel. The statistical significance is displayed as follows: *p < 0.05; **p < 0.01; ***p < 0.001; ****p < 0.0001; ns, not significant. Panels B-D: one-way ANOVA with FDR correction’ Panels E, G: Student’s t-test (Gaussian assumption). Panel H: Unpaired Welsh’s t-test, data plotted as geometric mean.

In addition to requiring aspartate, pyrimidine biosynthesis depends on mitochondrial respiration as dihydroorotate dehydrogenase (DHODH), a key pyrimidine biosynthetic enzyme, donates electrons to the electron transport chain (ETC)^45^ (Figure 7A). This dependence can be rescued via uridine supplementation, which supplies pyrimidines via the salvage pathway^46^. We therefore tested whether aspartate and uridine supplementation can rescue HCMV-mediated induction of dNTP levels. Supplementation with aspartate and uridine significantly increased several dNTP levels upon adaptaquin treatment during HCMV infection (Figure 7D). Further, aspartate and uridine supplementation largely rescued viral DNA replication upon adaptaquin treatment (Figure 7E). This increase in viral DNA replication correlated with substantially increased viral spread upon aspartate and uridine supplementation (Figure 7F&G), highlighting the importance of prolyl hydroxylase activity for dNTP production to support viral DNA replication and productive infection. Collectively, our results indicate that HCMV relies on EGLN activity to drive mitochondrial activation supporting the production of dNTP precursors necessary for robust infection.

### Adaptaquin inhibits HCMV replication *in vivo*

We next tested the efficacy of adaptaquin in vivo utilizing a humanized mouse model. After implantation of infected human fibroblasts, mice received daily intraperitoneal injections of adaptaquin for nine consecutive days. At the end of the treatment period, plugs were harvested and viral output was quantified. Adaptaquin treatment resulted in a significant reduction in viral progeny relative to vehicle-treated controls, with efficacy comparable to valganciclovir (VGCV) treatment, the standard of care antiviral for HCMV infection (Figure 7H). Additionally, adaptaquin administration did not increase most serum markers of hepatotoxicity assessed including aspartate aminotransferase, alkaline phosphatase, and bilirubin, although alanine aminotransferase was moderately increased with adaptaquin treatment (Supplemental Fig. 5). These results indicate that adaptaquin suppresses HCMV replication *in vivo,* suggesting that targeting EGLN activity could be a novel avenue for anti-viral development.

## Discussion

Viruses parasitize cellular metabolic resources to drive the mass production of viral progeny. Regulation of these resources has emerged as a critical host-pathogen interaction that can determine the outcome of viral infection. However, while viral infection has been shown to depend on core central carbon pathways, precise understanding of specific viral metabolic vulnerabilities remains limited. Furthermore, the specific mechanisms governing metabolic regulation during infection are largely unclear. To address these issues, we utilized a metabolism-focused pharmacological library to screen for HCMV-associated metabolic vulnerabilities. Our results indicate that HCMV replication depends on specific mitochondrial, nucleotide, amino acid, and lipid-associated metabolic activities (Figure 1). We focused on the EGLN family of prolyl hydroxylases, which we find is necessary for HCMV-mediated activation of respiration and productive infection. Specifically, EGLN1 is required during infection to increase mitochondrial respiration and produce the TCA cycle-associated metabolites necessary to support dNTP biosynthesis and viral DNA replication.

We developed an anti-HCMV screening pipeline that employs a compound library targeting enzymes associated with metabolism arrayed over a 10-point dose curve that has been utilized to examine metabolic vulnerabilities in various contexts^21,22^. This pipeline identified several novel compounds that inhibit HCMV infection (Supplemental File 1). Several compounds inhibited pathways previously identified to be important for HCMV infection, although many targeted novel enzymatic targets within these pathways. For example, inhibition of aspartate transcarbamylase, an important pyrimidine biosynthetic enzyme, has previously been shown to inhibit HCMV infection^23^. Here, we find that brequinar, an inhibitor of the downstream DHODH-catalyzed reaction, also attenuates HCMV replication (Figure 1C). In addition to identifying novel enzymatic targets in pathways reported to be important for HCMV replication, our screen highlighted novel putative metabolic dependencies, including phosphoglycerate dehydrogenase, a key serine biosynthetic enzyme, and 5-lipoxygenase, a leukotriene biosynthetic enzyme. These enzymes are inhibited by NCT-503 and CJ-13610, respectively, and both were among the top compounds to limit HCMV infection (Figure 1C).

We primarily focused on inhibitors of infection, but some compounds in the screen appeared to stimulate infection (Figure 1B). While these potential pro-viral effects require validation, the ability to mass-produce high-titer viral stocks for vaccine production or for gene therapy vector production remains a substantial pharmaceutical challenge^47^. Our screen suggests that metabolism might be an attractive target to improve the production of viral vaccines or viral vectors.

Our results indicate that EGLN family members impact HCMV infection in different ways. While knockdown of EGLN1 or EGLN2 expression attenuated infection, knockdown of EGLN3 moderately enhanced infection (Figure 4). As prolyl hydroxylases, EGLN family members can exhibit distinct substrate affinities and specificities with different downstream consequences^48^. EGLN3 has been found to stimulate apoptosis^49^, which could restrict HCMV replication. Additionally, EGLN3 prolyl hydroxylates acetyl-CoA carboxylases (ACC), which induces ACC degradation^50^. Acetyl-CoA carboxylases catalyze a rate-determining step in fatty acid biosynthesis and regulate fatty acid oxidation. We have previously found these enzymes to be important for HCMV replication^11^, and our current screen identified an additional ACC inhibitor that attenuates HCMV infection (Figure 1F). EGLN3-mediated prolyl hydroxylation and subsequent degradation of ACC enzymes would be predicted to limit infection. Either of these EGLN3-associated activities, i.e., stimulating apoptosis or inducing ACC degradation, could be responsible for the increased HCMV replication upon targeting EGLN3 expression.

The knockdown of either EGLN1 or EGLN2 attenuated HCMV infection, albeit to different extents. These differences likely reflect different substrate specificities between isoforms. EGLN2 can hydroxylate FOXO transcription factors, resulting in their destabilization^51^, whereas EGLN1 has been shown to hydroxylate AKT, resulting in its inhibition^52^, and to regulate RAF-ERK activity via hydroxylation of the c-RAF activator, NDRG3^53^. Notably, CRISPRi-mediated knockdown of the different enzymes substantially differed with respect to their impact on mitochondrial respiration (Figure 5). EGLN1 targeting largely ablated HCMV-mediated activation of respiration, whereas EGLN2 targeting had little impact, slightly increasing uncoupled respiration (Figure 5B&C). However, interpreting the contributions of the individual EGLNs to mitochondrial remodeling is complicated by the fact that modulation of the expression of one EGLN family member impacted the expression of the other EGLN isoforms (Supplemental Figure 3).

Over seventy years ago, influenza infection was found to induce cellular respiration, and further, that inhibitors of either the TCA cycle or oxidative phosphorylation inhibited influenza infection *in vitro* and *in vivo*^54,55^. Soon after, oxidative phosphorylation inhibitors were found to attenuate the replication of various evolutionarily diverse viruses^56^. The earliest assumption, which remains prevalent, is that viral infection induces TCA activity to provide the energy necessary to produce viral progeny. Our results indicate that viral infection depends on TCA cycle activation to provide the biosynthetic precursors necessary to support viral DNA replication.

Adaptaquin significantly inhibited the replication of a common cold coronavirus (OC43) (Figure 2K). However, it did not inhibit HSV-1 replication (Figure 2J), despite HSV-1 being a herpesvirus in the same family as HCMV. Notably, these viruses interact with cellular nucleic acids very differently. HSV degrades both host mRNA^57^ and mitochondrial DNA^58^, which could provide pools of nucleotides to support nucleic acid synthesis, and explain the lack of adaptaquin-mediated inhibition. In contrast, HCMV and OC43 do not degrade cellular nucleic acids, highlighting a potential increased reliance on the mitochondrial production of nucleotide precursors. Future research will need to explore the extent to which different viruses rely on EGLN activity to support productive infection.

EGLN1 has been reported to translocate to mitochondria in a HIF1α-independent manner^39^, suggesting that EGLN1 likely possesses mitochondria-associated functions independent of infection. Consistent with this, we find that adaptaquin treatment reduced mitochondrial biogenesis in uninfected cells (Figure 5F). However, to our knowledge, EGLN1 has not been shown to modulate respiration. Further, putative mitochondrial prolyl hydroxylation targets that could impact respiration or mitochondrial physiology remain to be elucidated. Given the important roles that the EGLN family plays in oncogenesis^59,60^, tissue repair^61^, and metabolism^62^, elucidating the targets through which EGLN activity could modulate mitochondrial function remains a key priority.

Collectively, our results indicate that EGLN1 is an important host factor for viral-mediated mitochondrial remodeling, driving the production of substrates required for viral DNA replication. Given that HCMV induces significant morbidity in neonates and immunosuppressed individuals^1^ and current treatment options exhibit significant limitations, e.g., the emergence of drug resistance, poor bioavailability, and toxicity^5,6^, novel therapeutic strategies are necessary. EGLN inhibitors are currently in various stages of clinical testing for a variety of purposes, including improving cardiovascular responses to ischemia^63^, promoting erythropoiesis to treat anemia^64,65^, and reducing the adverse effects of radiotherapy^60^. These trials suggest that EGLN inhibitors might be therapeutically useful, and accordingly, that their potential as anti-viral agents should be further explored.

## Materials and Methods

### Cell culture, viruses, and viral infections

Telomerase-immortalized MRC5 (MRC5-hT) lung fibroblasts and Human Foreskin Fibroblasts were cultured in Dulbecco’s modified Eagle serum (DMEM; Invitrogen #11965118) supplemented with 10% (v/v) fetal bovine serum (Biowest) and 1% penicillin-streptomycin (Pen-Strep; Life Technologies #15140-122) at 37°C in a 5% (v/v) CO_2_ atmosphere.

Unless indicated otherwise, wild-type viral infections were performed using a virus derived from a BADwt bacterial artificial chromosome (BAC) clone of a GFP-expressing, UL131Repair HCMV AD169 viral strain^66^. The TB40/E-UL99-mCherry virus (referred to as HCMV-mCherry) was a gift from Christine M. O’Connor and Eain Murphy, generated as described^67^. All viral stocks used in this manuscript were propagated in MRC5 cells. For experiments involving infection, MRC5 cells were grown to confluence and then placed in serum-free DMEM supplemented with 1% PenStrep for 24 hours prior to infection. The medium was replaced with viral adsorption medium to cover the cells containing the indicated MOI in serum-free DMEM. After 2 hour adsorption, the medium was removed and replaced with fresh serum-free DMEM with 1% PenStrep unless otherwise stated. For analysis of viral titers, infectious virus was quantified via serial dilution and SV40-promoter-associated GFP expression analysis in MRC5-hT cells at 48 hpi unless otherwise indicated. For HSV-YFP^68^ infections, MRC5-hT cells were grown to 100% confluence and subsequently infected in serum-free DMEM at an MOI of 0.05 (Low MOI) or 5 (High MOI). Virus-containing media was harvested at 24 hours post infection. For OC43 infections, MRC5-hT cells were grown to 100% confluence and subsequently infected in DMEM containing 2% FBS at an MOI of 0.05 (Low MOI) or 1 (High MOI). Virus-containing media was harvested at 24 hours post infection. HSV and OC43 virion production was assessed via TCID50/mL, which was quantified as previously described^69^.

### Compound library and high-throughput screening

The compound library for the high-throughput screen was prepared as described previously^21^. Before screening with the compound library, the Z′ factor was measured to determine the robustness of our assay^70^.

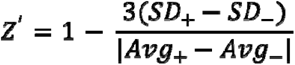

A 384-well plate (Corning #3764) was seeded with 6,000 cells in a volume of 50 μL using a Multidrop Combi reagent dispenser (Thermo Scientific). Cells were incubated at 37°C for 24 hours, following which the medium was exchanged with serum-free DMEM and incubated for an additional 24 hours. Following this, the media was exchanged again, with half of the wells receiving DMEM with added phosphonoacetic acid (PAA), a viral DNA polymerase inhibitor, and half receiving DMEM with added DMSO at the same volume of PAA. PAA was utilized as a positive control for viral inhibition due to its ability to inhibit viral DNA replication^71^. Cells were then incubated at 37°C for 9 days, whereupon they were fixed with 4% paraformaldehyde. GFP-positive cells were then quantified as described below, and the average and standard deviation of the number of infected cells were used to determine the Z’ factor of 0.955.

To determine compounds that inhibit HCMV replication, cells were seeded and serum-starved as described above. After switching to serum-free media, 100 nL of compounds from the library plates was pin transferred onto the cells and incubated at 37°C for 9 days, at which point they were fixed with 4% paraformaldehyde and nuclei stained with 1x Hoescht 33342 (Invitrogen, H3570). Cells were then imaged using a Cytation 5 automated imaging reader (Agilent, formerly known as BioTek). Each well was imaged using a 4x magnification objective lens, a predefined DAPI channel with an excitation wavelength of 377 nm and emission wavelength of 447 nm, and a predefined YFP channel with an excitation wavelength of 469 nm and emission wavelength of 525 nm. Gen5 3.11 software (Agilent, formerly known as BioTek) was used to determine cell number by gating for objects in both the DAPI and YFP channels. Data post-processing was conducted using an R script, followed by analysis in GraphPad Prism.

### Viral DNA accumulation

MRC5-hT cells were grown to confluence in a 12-well plate (Greiner) and infected as described above. Cells were harvested at the indicated time post-infection in 250 μL Lucigen QuickExtract DNA Extraction Solution, following the manufacturer’s protocol. DNA was diluted 1:100 in sterile H2O and analyzed by RT-qPCR. Viral DNA was quantified with IE1 probe and normalized to the background signal associated with mock-infected cells.

### Mitochondrial DNA accumulation

MRC5-hT cells were grown to confluence in a 12-well plate (Greiner) and infected as described above. Cells were harvested at the indicated time post-infection in 250 μL Lucigen QuickExtract DNA Extraction Solution, following the manufacturer’s protocol. DNA was diluted 1:100 in sterile H2O and analyzed by RT-qPCR using ND1 and B-actin primers described in the Real Time qPCR section of these methods. Quantities of ND1 compared to B-actin were calculated as previously described^42^.

### Cellular Fractionation

MRC5-hT cells were grown to confluence in T25 cell culture flasks (Greiner) and infected as described above. Cells were harvested at the indicated time post-infection, and cell fractions were isolated utilizing the Cell Fractionation Kit (Cell Signaling #9038). 2% of total starting cell equivalents were loaded for subsequent western blotting analysis.

### Mitochondrial respiration assays

MRC5-hT cells were grown to confluence in Seahorse XFe96 cell culture microplates (Agilent, Seahorse FluxPak, 103793-100) and infected as described above. One day prior to the assay, a Seahorse XFe96 Sensor cartridge was hydrated with XF calibrant (Agilent) in a hydrated, non-CO_2_ 37°C incubator. The day of the assay, the culture medium was removed and replaced with Agilent Seahorse XF DMEM medium (Agilent 103474-100) supplemented with 1mM GlutaMax (Gibco 35050-061), 1mM Sodium Pyruvate (Gibco 11360-070), and 10mM Glucose Solution (Gibco A2494001). Mitochondrial function was assessed using the Seahorse XF Cell Mito Stress Test (Agilent 103015-100) on a Seahorse XF96 analyzer. Oxygen consumption rate (OCR) was normalized to cell number as determined by Hoescht nuclear staining, as described above.

### Lentiviral transduction and cell line generation

To generate CRISPRi EGLN constructs, guide RNAs targeting EGLN1, EGLN2, and EGLN3 with flanking sequences (listed below) to the CRISPRi Lentivirus (LV) PURO plasmid (pLV hU6-sgRNA hUbC-dCas9-KRAB-558 T2a- puro with stuffer), gifted by the Harris Lab at the University of Rochester Department of Biomedical Genetics.

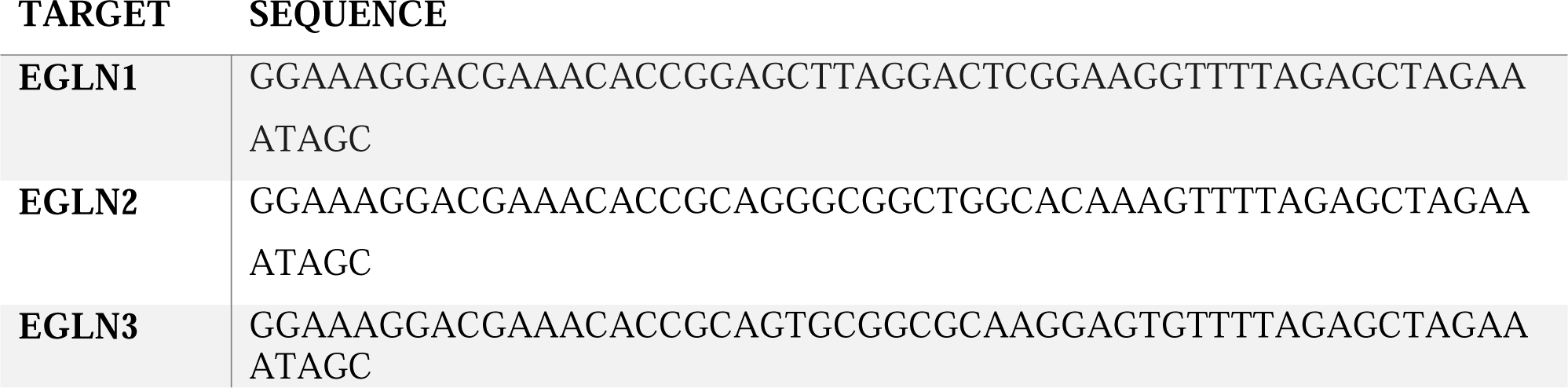

Lentivirus was generated in 50-70% confluence HEK 293T cells. PBS containing psPAX2 (Addgene #12260), pMD2.G (Addgene #12259), and our lentiviral plasmid was incubated for 5 minutes. Following this incubation, FuGENE (Promega, E2691) was added to the mixture and incubated for 20 minutes. This solution was added to HEK 293T cells and incubated at 37°C overnight. The transfection medium was then aspirated and replaced with DMEM with 10% FBS and 1% penicillin-streptomycin. Plates were incubated overnight, and then the medium was harvested by filtration with a 45 µm syringe filter. Lentiviral media were flash frozen and stored at -80°C. MRC5-hT cells were grown to 50-70% confluence, and the media were replaced with lentiviral inoculum with 8μg/mL polybrene. Lentiviral inoculum was replaced 3x after 6 hours incubation for a triple transduction with lentivirus prior to antibiotic selection in 1ug.mL puromycin containing medium. Antibiotic selection was performed until a non-transduced control place of MRC5-hT cells had died.

### Matrigel-fibroblast model for anti-HCMV activity in vivo

MRC5-hT cells were infected with TB40/E-UL99-mCherry at an MOI of 0.02 in DMEM with 10% FBS and 1% penicillin-streptomycin. Cells were incubated overnight, and then the cells were resuspended in 1 mL of PBS and added to a Matrigel (Corning #354262) suspension at 4 x 10^6^ cells per mL. 400 uL (1.6 x 10^6^) matrigel suspension was loaded into syringes and placed on ice. NOD.Cg-Prkdcscid II2rgtm1Sug/JicTac mice (CIEA NOG, model no. NOG-F, genotype sp/sp;ko/ko, Taconic Biosciences) were anesthetized using isoflurane, the dorsolateral region above the left hip was shaved, and the region was sterilized with Betadine (Henry Schein #6906950) and 70% Isopropanol. Matrigel was injected subcutaneously into the left flank. Adaptaquin (50 mg/kg, i.p., daily), valganciclovir (200 mg/kg, i.p., daily), or vehicle control was administered, beginning at the time of implantation. On day 9, mice were euthanized and the matrigel plugs were harvested. Matrigel plugs were digested with 0.5 mg/mL Collagenase Type II (Sigma-Aldrich, C2-28-100MG) for 3 hours with vortexing every hour. The resulting suspensions were flash frozen, thawed, and sonicated to release infectious virus. Suspensions were clarified by centrifugation at 2,000 rpm for 5 minutes, and the supernatant was passed through a 40-µm cell strainer (Greiner, 76469-404). Infectious units were determined as described above and plotted as geometric means. Adaptaquin was prepared as a 4 mg/mL suspension in 5% DMSO, 40% PEG300 (Thermo Scientific AC192220010), 5% Tween 80 (Thermo Scientific AC278632500), 50% ddH20. Valganciclovir was prepared as a 16 mg/mL suspension in 98% PBS, 2% DMSO. A solution of 5% DMSO, 40% PEG300, 5% Tween 80, 50% ddH20 was used as a vehicle control.

### Cell viability assays

MRC5-hT cells were grown to confluence in a 96-well plate and treated with Adaptaquin for 5 days post-infection. One row of the cells was fixed with methanol as a positive control for apoptosis. Cells were either stained with 1x Hoescht 33342 (Invitrogen, H3570) and 500nM Propidium Iodide (MedChemExpress, HY-D0815) by incubating at 37°C for 60 minutes. Cells were then imaged via a Cytation 5 microscope.

### Real-time qPCR

RNA was isolated from cells using Trizol extraction followed by Direct-zol RNA Miniprep Plus kit (Zymo Research). Isolated RNA was used to synthesize cDNA with a qScript cDNA synthesis kit (Quantabio). Samples were analyzed by Real-Time qPCR, and relative quantities of gene expression were calculated as previously described^22^. The following primers were used:

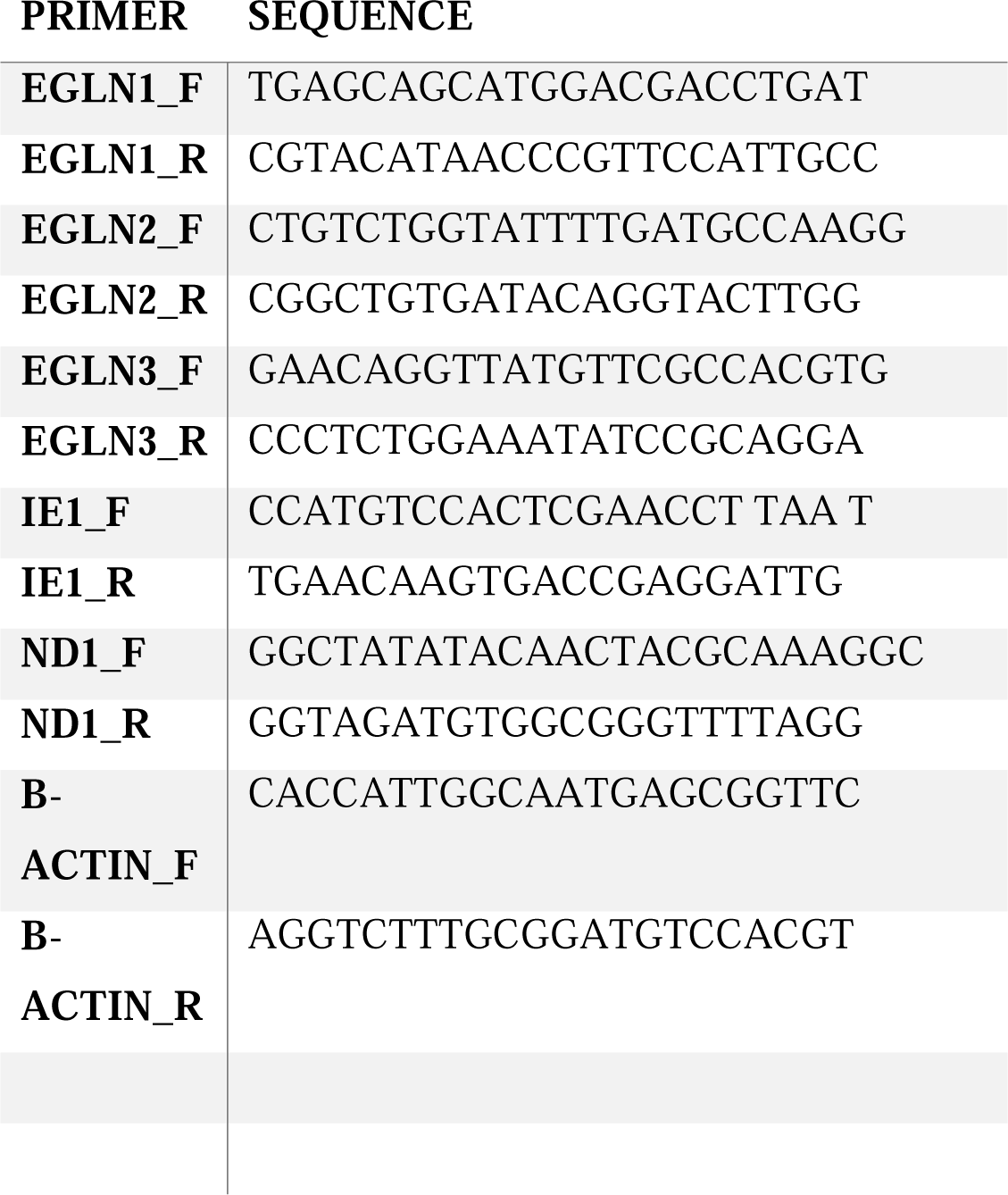

### Protein analysis and western blotting

Protein detection was performed as previously described^72^. The following antibodies were used for western blot analysis following the manufacturer’s instructions: Vinculin (Cell Signaling, 13901), PHD3/EGLN3 (Proteintech, 18325), Monoclonal Anti-FLAG (Sigma-Aldrich, F1804), IE1^73^, PP28^74^, UL26^75^.

### Proteomics

Cells were lysed in 200 μl of 5% SDS, 100 mM TEAB and sonicated (QSonica) for 5 cycles, with a 1-minute resting period on ice after each cycle. Samples were then centrifuged at 15,000 x g for 5 minutes to pellet cellular debris, and the supernatant was collected. Protein concentration was determined by BCA (Thermo Scientific), after which samples were diluted to 1 mg/mL in 5% SDS, 50 mM TEAB. 25 µg of protein from each sample was reduced with dithiothreitol to 2 mM, followed by incubation at 55°C for 60 minutes. Iodoacetamide was added to 10 mM and incubated in the dark at room temperature for 30 minutes to alkylate the proteins.

Phosphoric acid was added to 1.2%, followed by six volumes of 90% methanol, 100 mM TEAB. The resulting solution was added to S-Trap micros (Protifi) and centrifuged at 4,000 x g for 1 minute. The S-Traps containing trapped protein were washed twice by centrifuging through 90% methanol, 100 mM TEAB. 1 µg of trypsin was brought up in 20 µL of 100 mM TEAB and added to the S-Trap, followed by an additional 20 µL of TEAB to ensure the sample did not dry out. Samples were incubated at 37°C overnight. The next morning, the S-Trap was centrifuged at 4,000 x g for 1 minute to collect the digested peptides. Sequential additions of 0.1% TFA in acetonitrile and 0.1% TFA in 50% acetonitrile were added to the S-trap, centrifuged, and pooled. Samples were frozen and dried down in a Speed Vac (Labconco), then re-suspended in 0.1% trifluoroacetic acid prior to mass spectrometry analysis. Peptides were injected onto a homemade 30 cm C18 column with 1.8 μm beads (Sepax), with an Easy nLC-1200 HPLC (Thermo Fisher), connected to a Fusion Lumos Tribrid mass spectrometer (Thermo Fisher). Solvent A was 0.1% formic acid in water, while solvent B was 0.1% formic acid in 80% acetonitrile. Ions were introduced to the mass spectrometer using a Nanospray Flex source operating at 2 kV. The gradient began at 3% B and held for 2 minutes, increased to 10% B over 5 minutes, increased to 38% B over 68 minutes, then ramped up to 90% B over 3 minutes and was held for 3 minutes, before returning to starting conditions over 2 minutes and re-equilibrating for 7 minutes, for a total run time of 90 minutes. The Fusion Lumos was operated in data-dependent mode, with MS1 and MS2 scans acquired in the Orbitrap. The cycle time was set to 2 seconds. Monoisotopic Precursor Selection (MIPS) was set to ‘Peptide’. The full scan was performed over a range of 375-1400 *m/z*, with a resolution of 120,000 at *m/z* of 200, an AGC target of 4e5, and a maximum injection time of 50 ms. Peptides with a charge state between 2-5 were picked for fragmentation, with minimum and maximum intensity thresholds set to 1e5 and 1e20, respectively. Precursor ions were fragmented by higher energy collision dissociation (HCD) using a collision energy of 30% with an isolation width of 1.1 *m/z*. The fragment scan was performed with a resolution of 15,000 at *m/z* of 200, an AGC target of 5e4, and a maximum injection time of 25 ms. Dynamic exclusion was set to 20 seconds with a mass tolerance of 10 ppm.

Raw data were searched using the SEQUEST search engine within the Proteome Discoverer software platform, version 2.4 (Thermo Fisher), using the SwissProt human and HCMV databases. Trypsin was selected as the enzyme, allowing up to 2 missed cleavages, with an MS1 mass tolerance of 10 ppm, and an MS2 mass tolerance of 0.6 Da. Carbamidomethyl was set as a fixed modification, while oxidation of methionine and proline was set as variable modifications. The Minora node was used to determine relative protein abundance between samples using the default settings. Percolator was used as the FDR calculator, filtering out peptides that had a q-value greater than 0.01.

### Metabolomics Sample Preparation, LC-MS Acquisition, and Data Processing

MRC5-hT cells were grown to confluence in 6-well plates. Medium was replaced with serum-free DMEM for 24 h prior to infection. At 48 h post-infection, 2 h before harvest, medium was replaced with serum-free DMEM pre-equilibrated at 37 °C in a 5% CO atmosphere. Plates were washed once with ice-cold PBS + glucose (Sigma G-8270; volume equal to previous medium), and 1 mL dry ice–cooled extraction solution was added to each well. Cells were scraped, pipetted up and down three times, and transferred to 1.5 mL tubes. Tubes were incubated for 15 min on dry ice, then for 30 min on ice, vortexing for 5 s every 10 min (three total vortex steps). Lysates were centrifuged at 14,000 rpm (max speed) for 10 min at 4 °C. Supernatant (900 µL) was collected, leaving ∼100 µL to avoid pellet disruption, dried under nitrogen, and stored at −80 °C.

Dried extracts were resuspended in 90 µL ice-cold 50% acetonitrile, vortexed 5 s, and incubated on ice for 40 min with vortexing for 5 s every 10 min (four total vortex steps). Samples were centrifuged at 14,000 rpm for 20 min at 4 °C. Aliquots (20 µL) were transferred to LC-MS vials; 8 µL from each sample was combined into a pooled quality control (QC) tube.

Metabolites were analyzed on an Orbitrap Exploris 240 (Thermo) coupled to a Vanquish Flex LC system (Thermo). Injections (2 µL) were made onto a Waters XBridge XP BEH Amide column (150 mm × 2.1 mm, 2.5 µm particle size) with guard column, held at 25 °C. For positive ion mode, mobile phase A was H O + 10 mM ammonium formate + 0.125% formic acid; mobile phase B was 90% acetonitrile + 10 mM ammonium formate + 0.125% formic acid. For negative ion mode, mobile phase A was H O + 10 mM ammonium acetate + 0.1% ammonium hydroxide + 0.1% medronic acid; mobile phase B was identical to positive mode. The gradient (150 µL min ¹ unless otherwise specified) was: 0–2 min, 100% B; 2–3 min, 90% B; 3–5 min, 90% B; 5–6 min, 85% B; 6–8 min, 75% B; 8–10 min, 55% B; 10–13 min, 35% B; hold to 20 min; 20.1–20.6 min, 100% B; 20.6–22.2 min, 100% B; 22.7–27.9 min, 100% B at 300 µL min ¹; 28 min, 100% B at 150 µL min ¹.

The H-ESI source was operated in positive mode at 3,500 V or negative mode at 2,500 V with sheath gas 35 au, auxiliary gas 7 au, sweep gas 0 au, ion transfer tube 320 °C, vaporizer 275 °C. Full-scan data were acquired from 70–1,000 m/z at 120,000 FWHM resolution, RF lens 70%, standard AGC.

LC-MS data were processed in Compound Discoverer v3.3 (Thermo) and El-MAVEN. Chromatography features from all uploaded files were aligned to a reference sample consisting of a pool of all of the other samples and that was run alongside the other samples using the ChromAlign algorithm^76^. In Compound Discoverer, features were aligned to pooled QC (3 ppm mass tolerance, 0.2 min RT tolerance), detected at ≥30,000 intensity, S/N ≥1.5, peak rating ≥3, and gap-filled (3 ppm, S/N ≥1.5, ≥3 scans). Features with sample:blank ratios ≤2 were removed. Annotation used mzVault, MassList, mzCloud, Predicted Compositions, ChemSpider, and Metabolika, with in-house RT-matched standards (RT tolerance 0.3–0.6 min). mzVault matches required score ≥90; mzCloud matches required confidence ≥50. In El-MAVEN, mzML files (MSConvertGUI, vendor peak picking, MS¹ only) were matched to curated positive (n=284) and negative (n=366) mode standard lists (3 ppm EIC tolerance, ±0.3 min RT) with filters: area ≥100,000, quality ≥0.70, signal/blank ≥2, signal/baseline ≥2, ≥3 scans peak width.

Positive and negative mode feature tables were merged in R. Duplicate entries (KEGG ID/CSID or name) were resolved by prioritizing: CD-mzVault > CD-MassList > El-MAVEN > CD-mzCloud > CD-ChemSpider/Metabolika > CD-Predicted Compositions. Known misannotations were corrected using a curated list. Annotation confidence tiers were: Tier 1a (mzVault ≥90 or MassList, unique mass/RT), Tier 1b (lower score or ambiguous, mzCloud ≥90/confidence ≥50), Tier 2 (mzCloud identified, but below Tier 1b 90/50 thresholds), Tier 3 (annotated MS1 match only (3ppm)), None (unannotated). High-confidence data included Tier 1a and Tier 1b with RT ≥5 min. For plotting in figures 6-7, cell-count-normalized metabolite peak areas were normalized by the maximum value for each metabolite across the samples and subsequently scaled to 100.

### Statistical analysis

Statistics were analyzed using GraphPad Prism v10 unless otherwise stated. Statistical comparisons were made using unpaired two-tailed t tests or ANOVA analysis with a normal (Gaussian) distribution assumption as indicated. For metabolomics data, cell normalized peak areas were auto scaled prior to one-way ANOVA analysis with FDR correction (Banjamini-Hochburg procedure) and subsequent Fisher’s LSD post-hoc testing as implemented in MetaboAnalyst^77^. Principle Component Analysis of metabolite data was performed in Metaboanalyst, with 95% confidence regions represented via shaded ellipse. For comparison of curve areas, e.g., with viral spread experiments, significance was assessed using 95% confidence intervals calculated in Prism according to the Gagnon method. For in vivo viral replication data, comparisons were made using an unpaired Welsh’s ttest.

## Acknowledgements

The work described in this publication benefited from the support of the Metabolomics Shared Resource at the Wilmot Cancer Institute, supported in part by the University of Rochester Wilmot Cancer Institute Support Grant P30CA272302. Further, the proteomics experiments were performed by the Mass Spectrometry Resource Lab at the University of Rochester Medical center. LAS was supported by an NIH T32 training grant (AI118689), as was J.C. who was supported by AI049815 and GM068411. The work was also supported by the following NIH grants to JCM; AI150698, AI181865 AI184380. The content is solely the responsibility of the authors and does not necessarily represent the official views of the National Institutes of Health. We would also like to thank Emily Tuttle for her guidance and advice on our in vivo experiments.

---

**Supplemental Figure 1.**
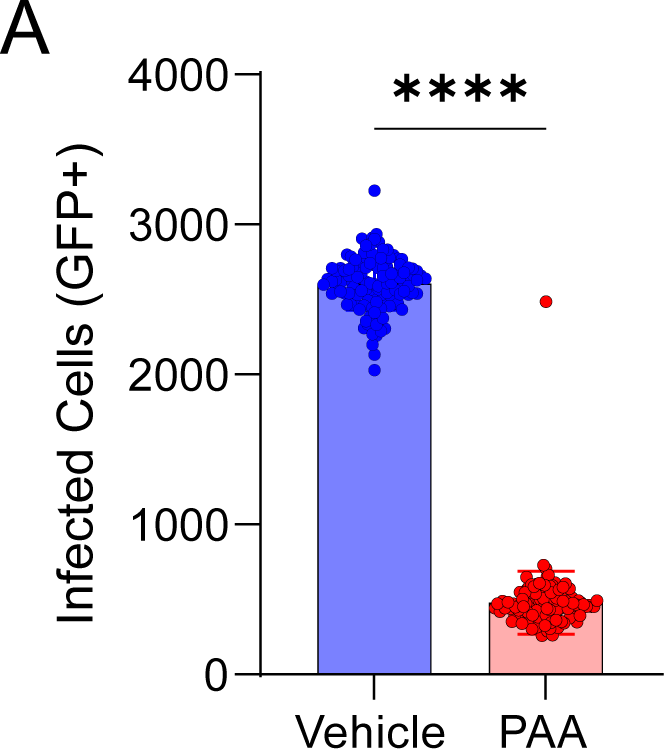
Z’ calculations confirm that the established screening parameters are suitable for a replicable high-throughput screen. (A) MRC5-hT cells were seeded in a 384 well plate and infected (MOI = 0.01). Cells were treated with PAA [50 ug/mL] or equal volume DMSO 2 hours post infection. Cells were fixed and imaged for GFP positivity 9 days post infection.

**Supplemental Figure 2.**
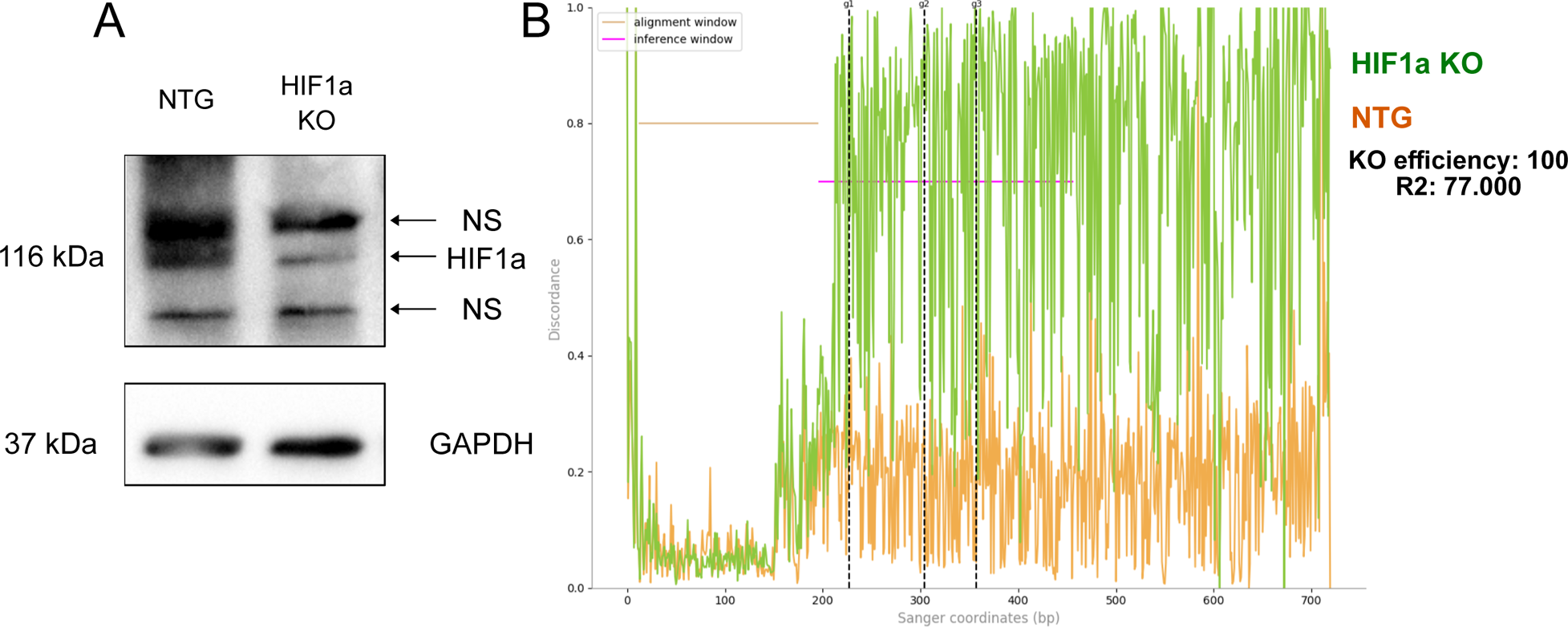
Genotyping and protein analysis confirm HIF1a knockout. (A) HFF cells were genome edited with ribonucleoprotein-mediated CRISPR. Proteins were harvested for western blot analysis. (B) Discordance plot from Synthego ICE analysis of HIF1a-knockout cells. On the plot, the level of alignment from the knockout cells (green) and control cells (orange) diverges following the cut site (dotted black lines).

**Supplemental Figure 3.**
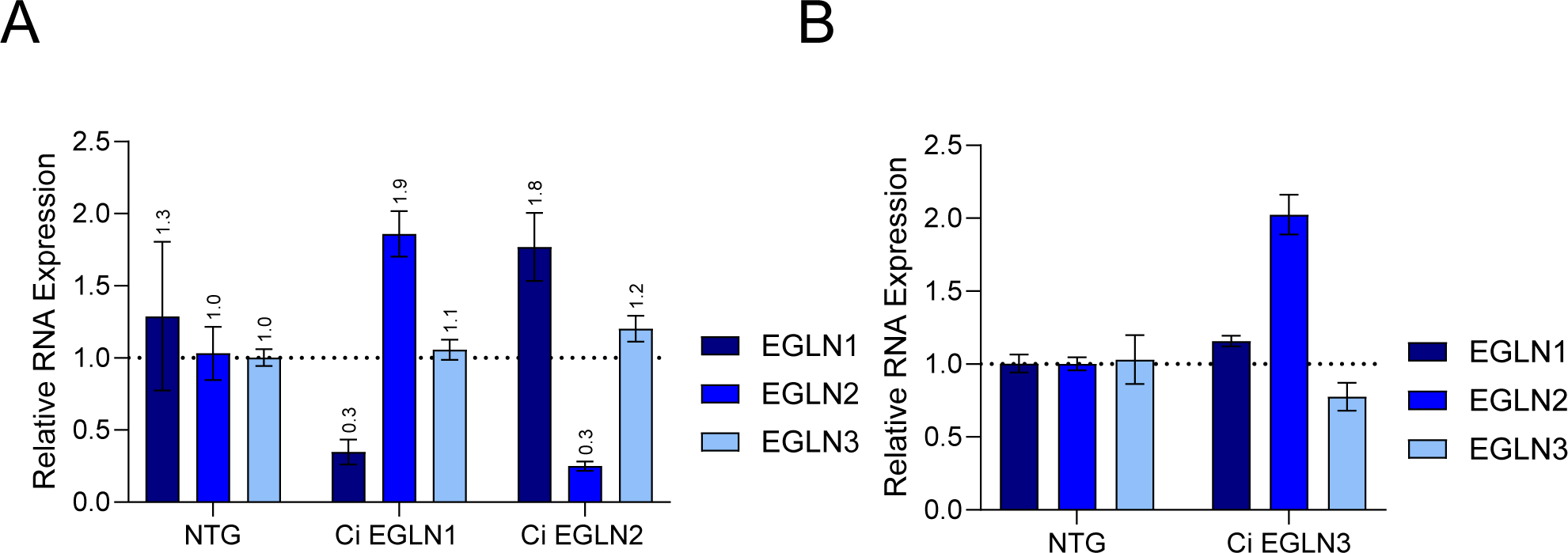
RNA expression in CRISPRi-EGLN constructs show compensation from other genes. (A) MRC5-hT cells were lentivirally transduced with a CRISPRi-construct targeting EGLN1 and EGLN2, respectively. RNA was harvested 48 hours post infection and EGLN expression assessed. (B) MRC5-hT cells were lentivirally transduced with a CRISPRi-construct targeting EGLN3. RNA was harvested 48 hours post infection and EGLN expression assessed.

**Supplemental Figure 4.**
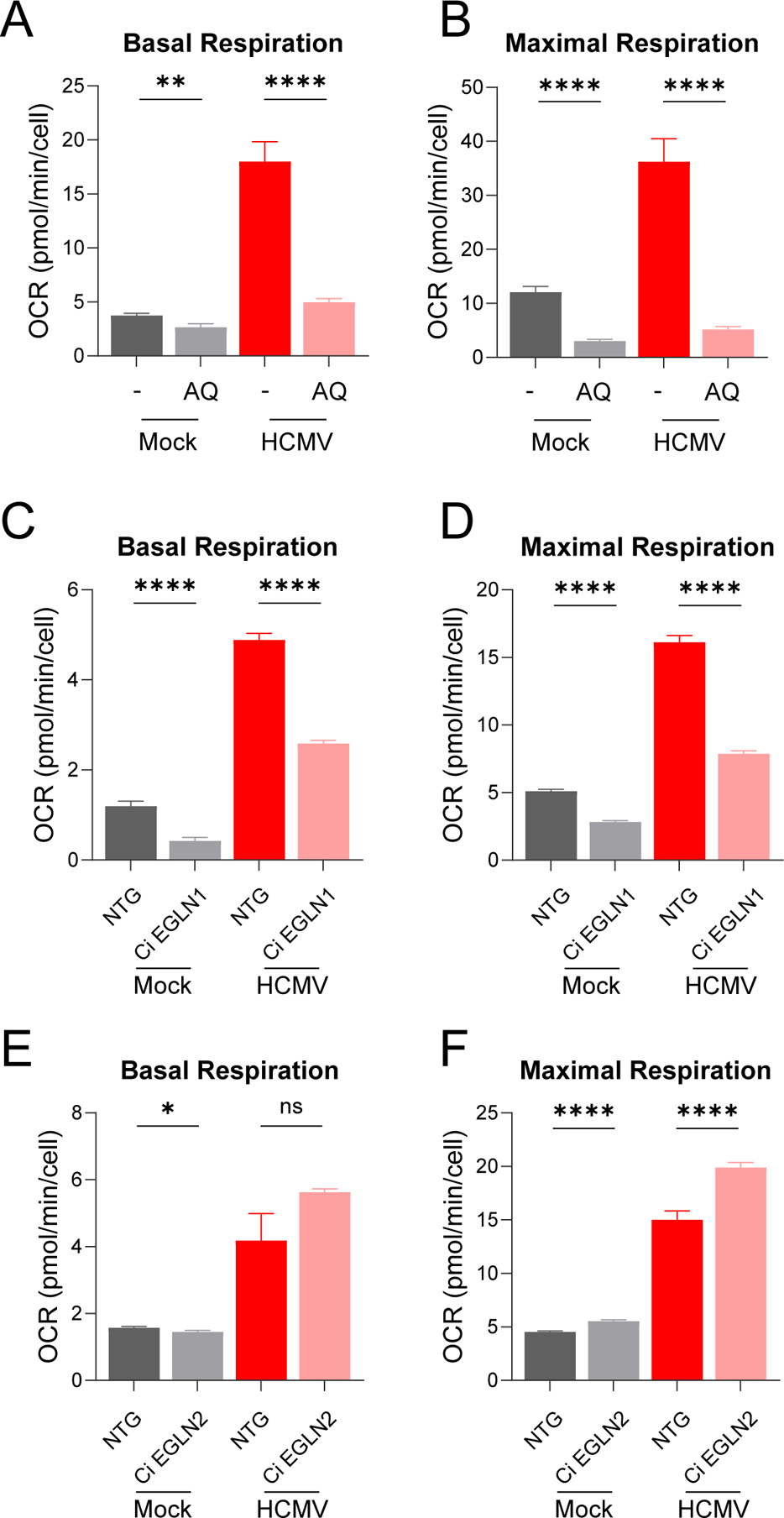
EGLN inhibition modulates mitochondrial basal and maximal respiration. (A-B) Basal and maximal respiration for adaptaquin (AQ) treated cells. Oxygen consumption normalized to cell count (MOI=1.5, Mean ± SEM, N=20). (C-D) Basal and maximal respiration for EGLN1 knockdown cells. Oxygen consumption normalized to cell count (MOI=1.5, Mean ± SEM, N=20). (E-F) Basal and maximal respiration for EGLN2 knockdown cells. Oxygen consumption normalized to cell count (MOI=1.5, Mean ± SEM, N=20). The statistical significance is displayed as follows: *p < 0.05; **p < 0.01; ***p < 0.001; ****p < 0.0001; ns, not significant.

**Supplemental Figure 5.**
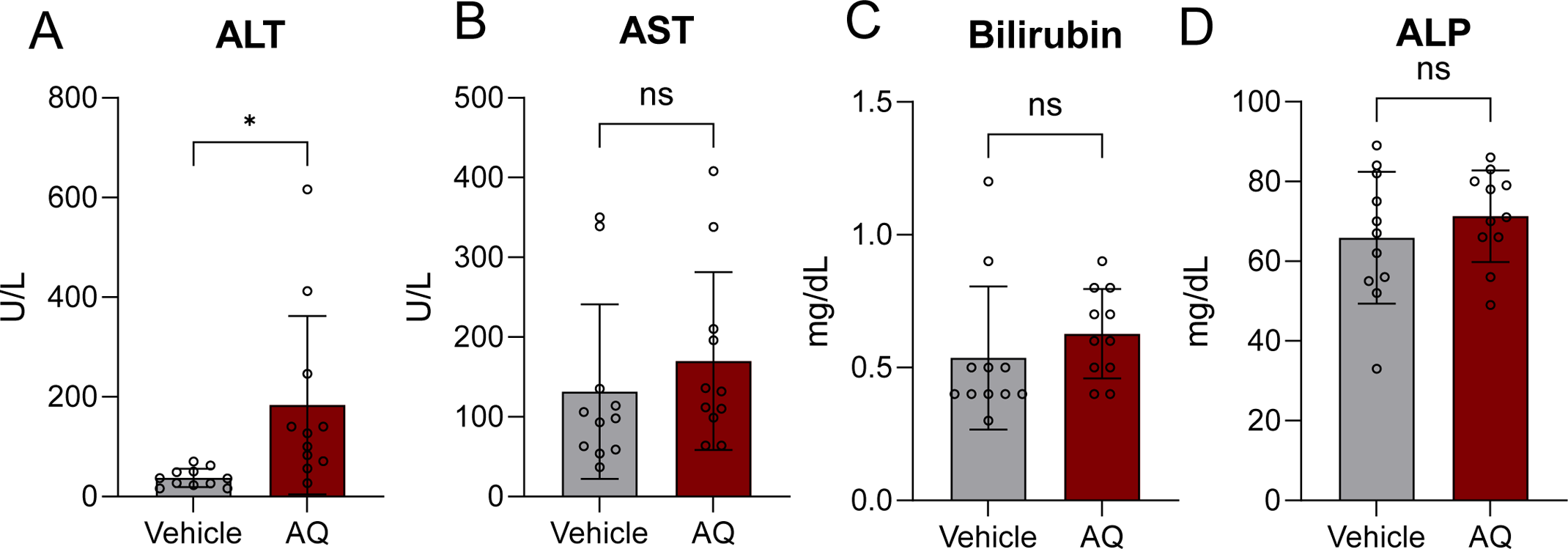
Adaptaquin induces a modest ALT increase without broader liver marker changes. Serum markers of liver function were measured following Adaptaquin treatment. (A) ALT levels were increased compared with vehicle controls. (B) AST, (C) total bilirubin, and (D) alkaline phosphatase (ALP) remain unchanged. The statistical significance is displayed as follows: *p < 0.05; ns, not significant.

